# Calcium-Binding Proteins are Altered in the Cerebellum in Schizophrenia

**DOI:** 10.1101/2020.03.02.972810

**Authors:** Francisco Vidal-Domènech, Gemma Riquelme, Raquel Pinacho, Ricard Rodriguez-Mias, América Vera, Alfonso Monje, Isidre Ferrer, Luis F. Callado, J. Javier Meana, Judit Villén, Belén Ramos

**Affiliations:** Psiquiatria Molecular, Institut de Recerca Sant Joan de Déu, Santa Rosa 39-57, 08950, Esplugues de Llobregat, Spain; Dept. de Bioquímica i Biologia Molecular, Facultat de Medicina, Universitat Autònoma de Barcelona, Bellaterra, 08193, Spain; Department of Genome Sciences, School of Medicine. University of Washington, Seattle, USA; Parc Sanitari Sant Joan de Déu, Dr. Antoni Pujadas, 42, 08830 Sant Boi de Llobregat, Spain; Departamento de Patologia y Terapeutica Experimental, Universidad de Barcelona; Senior consultant Servicio Anatomia Patológica, Hospital Universitario de Bellvitge-IDIBELL; CIBERNED; Hospital de Llobregat, Barcelona, Spain; Departament of Pharmacology, University of the Basque Country UPV/EHU, 48940 Leioa, Bizkaia, Spain; Centro de Investigación Biomédica en Red de Salud Mental, CIBERSAM, Spain; Biocruces Bizkaia Health Research Institute, Spain

## Abstract

Alterations in the cortico-cerebellar-thalamic-cortical circuit might underlie the diversity of symptoms in schizophrenia. However, molecular changes in cerebellar neuronal circuits, part of this network, have not yet been fully determined. Using LC-MS/MS, we screened altered candidates in pooled grey matter of cerebellum from schizophrenia subjects who committed suicide (n=4) and healthy individuals (n=4). Further validation by immunoblotting of three selected candidates was performed in two cohorts comprising schizophrenia (n=20), non-schizophrenia suicide (n=6) and healthy controls (n=21). We found 99 significantly altered proteins, 31 of them previously reported in other brain areas by proteomic studies. Transport function was the most enriched category, while cell communication was the most prevalent function. For validation, we selected the vacuolar proton pump subunit 1 (VPP1), from transport, and two EF-hand calcium-binding proteins, calmodulin and parvalbumin from cell communication. All candidates showed significant changes in schizophrenia (n=7) compared to controls (n=7). VPP1 was altered in the non-schizophrenia suicide group and increased levels of parvalbumin were linked to antipsychotics. Further validation in an independent cohort of non-suicidal chronic schizophrenia subjects (n=13) and non-psychiatric controls (n=14) showed that parvalbumin was increased while calmodulin was decrease in schizophrenia. Our findings provide evidence of an dysregulation of calcium-binding proteins in the cerebellum in schizophrenia, suggesting an impact on normal calcium-dependent synaptic functioning of cerebellar circuits. Our study also links VPP1 to suicide behaviours, suggesting a possible impairment in vesicle neurotransmitter refilling and release in these phenotypes.

## Introduction

Schizophrenia constitutes a complex disorder with a mixture of symptoms and cognitive deficits. It is considered a brain disorder in which a malfunction of multiple regions in distributed brain macro- and microcircuits could lead to the different symptoms. According to the hypothesis of cognitive dysmetria based on neuroimaging findings, the alteration of specific components of the cortico-cerebellar-thalamic-cortical circuit could contribute to the impairment in mental coordination processes and lead to the emergence of symptoms in schizophrenia [1–3]. The cerebellum, as part of this circuit, has been suggested to play a role in the production of the diversity of symptoms and cognitive impairments in this disorder. Much evidence has been collected over the last few decades that supports this view of the involvement of the cerebellum in higher cognitive functions and in the pathophysiology of this neurodevelopmental disorder [4–6]. Based on the clinic signs of individuals with cerebellar lesions and functional neuroimaging studies in patients with schizophrenia, it has been suggested that the impairment of the cerebellar function could affect many altered domains in schizophrenia, including executive functions, working memory, language, attention, social cognition and emotion (reviewed in [4,6,7]). The lateral cerebellar cortex is involved in some of these cognitive abilities, such as working memory, executive functions and speech [5,8–10]. Molecular hypothesis-driven studies have found genes from the GABAergic and glutamatergic neurotransmission system with altered expression in the cerebellar cortex, supporting the idea that the dysregulation of these systems is also disrupted in the cerebellum [11–16]. Gene expression changes have been detected in the cerebellum using transcriptomic screenings in a few studies in schizophrenia [17–19]. However, in addition to the critical role the cerebellum could play in this disorder, to the best of our knowledge proteomic strategies to identify meaningful altered proteins in cerebellar neuronal circuits have not yet been performed.

Here, we designed a quick exploratory screening using quantitative proteomic analysis in pooled grey matter of the cerebellum from schizophrenia (1 pool of 4 samples) and control healthy individuals (1 pool of 4 samples) with the aim of discovering and validating consistently altered proteins in schizophrenia (SZ) (S1A Fig). A total of 47 subjects were used in this study for validation by immunoblot, first in the same samples of the proteomic analysis and then in a pilot extended cohort (control, n=7; SZ, n=7), which also included a non-schizophrenia suicide group (n=6) and also in an independent and larger cohort with non-suicidal chronic schizophrenia subjects (control, n=14; SZ, n=13) (S1A Fig). Our validation focused on three selected candidates with a possible relevant role in the disorder and detected with robust changes in the screening.

## Materials and methods

### Brain tissue samples

For proteomic analysis, we used post-mortem human brain tissue obtained from the *UPV/EHU brain collection,* from the cerebellum of subjects with schizophrenia who had committed suicide (n=4) and control subjects who had died in a traffic accident and who had had no history of psychiatric episodes (n=4) (Table 1 and S1 Table). Samples were obtained at autopsies in the Basque Institute of Legal Medicine, Bilbao, Spain, in compliance with policies of research and ethical boards for post-mortem studies. Toxicological screening for antipsychotics, antidepressants, and other drugs was performed at the National Institute of Toxicology, Madrid, Spain All deaths were subjected to retrospective analysis for previous medical diagnosis. Subjects with ante-mortem criteria for paranoid schizophrenia according to the Diagnostic and Statistical Manual of Mental Disorders (DSM-IIIR and DSM-IV) were matched to control subjects who had died from accidental causes in a paired design, based on gender, age, and post-mortem delay (PMD).

**Table 1:**
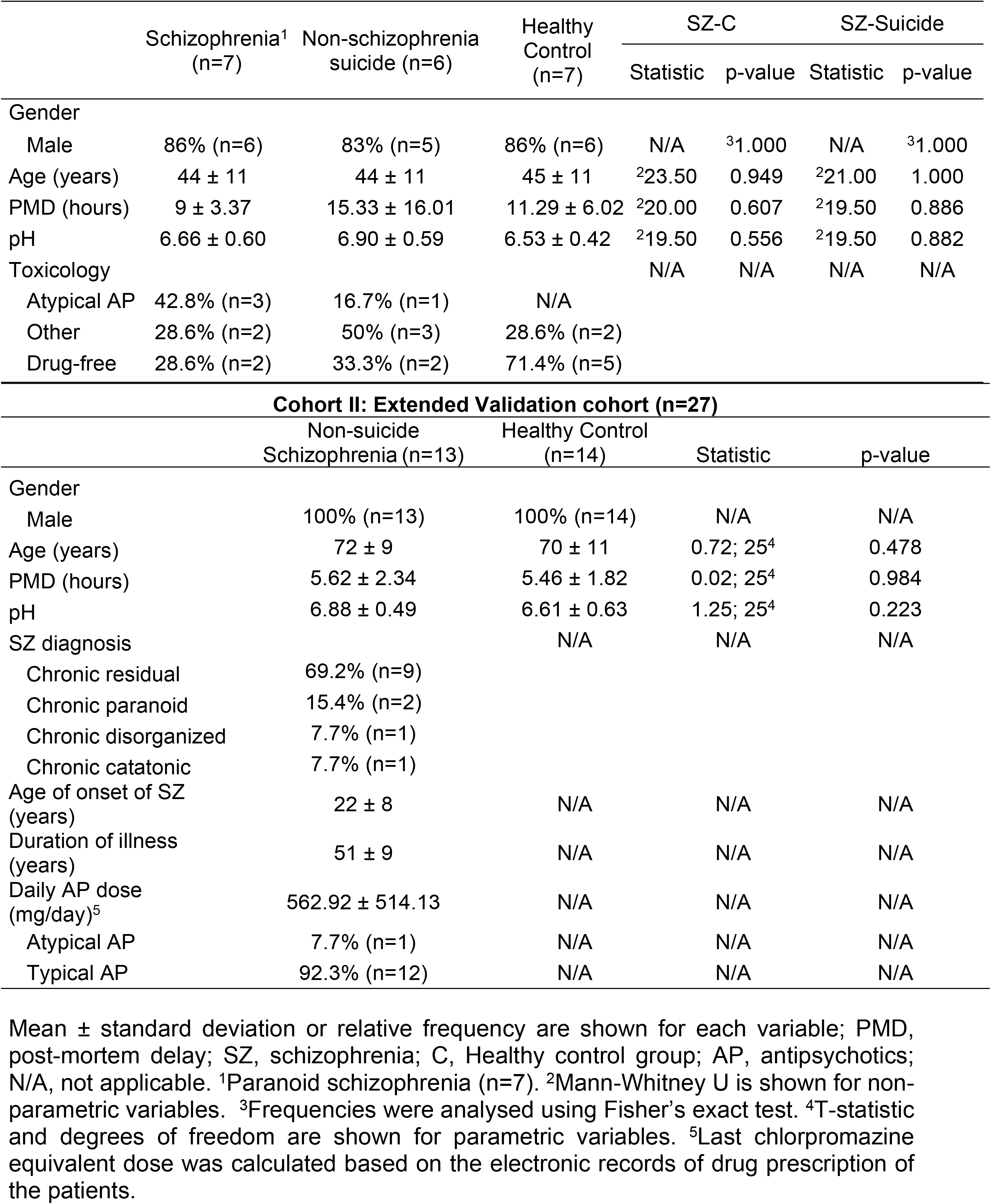
Demographic, clinical and tissue-related features of cases of Cohorts I and II.

To validate the candidates identified in the quantitative proteomic assay, we also used an extended cohort from the *UPV/EHU brain collection* for individual sample analysis of human post-mortem cerebellum (Table 1, Cohort I). A total of 20 brains of subjects who had committed suicide (n=11), died in an accident (n=5), through homicide (n=1) or from natural causes (n=3) were selected. Subjects with paranoid schizophrenia (n=7; 5 suicide victims and 2 non-suicide subjects), control subjects (n=7) that had died in an accident (n=5) or from natural causes (n=2), and a non-schizophrenia suicide victim group (n=6) were matched by gender, age, post-mortem delay and pH (Table 1, Cohort I). Control subjects were chosen among the collected brains on the basis, whenever possible, of the following criteria: (a) negative medical information on the presence of neuropsychiatric disorders or drug abuse, (b) accidental or natural cause of death, (c) negative results in toxicological screening for psychotropic drugs except for ethanol, and (d) a post-mortem delay not longer than 48 hours. Diagnoses were established according to the DSM-IIIR or DSM-IV. Diagnoses in the non-schizophrenia suicide victim group included adjustment disorder (n=3), anxiety disorder (n=1), personality disorder (n=1), and alcohol dependence (n=1). Samples were coded by the brain collection staff to protect human subject confidentiality. The study was approved by the Institutional Ethics Committee of the Fundació Sant Joan de Déu.

For further validation analysis of protein candidates, we also used an independent and larger cohort of post-mortem human cerebellum of subjects with chronic schizophrenia (n=13) that died by natural causes and control subjects with no history of psychiatric episodes (n=14) from the collection of neurologic tissues of Parc Sanitari Sant Joan de Déu [20] and the Institute of Neuropathology Brain Bank (HUB-ICO-IDIBELL Biobank) (Table 1, Cohort II) following the guidelines of Spanish legislation and the approval of the local ethics committees. Written informed consent was obtained from each subject. The study was approved by the Institutional Ethics Committee of Parc Sanitari Sant Joan de Déu. We matched schizophrenia and control groups by gender, age, post-mortem delay and pH Table 1 (Cohort II) shows the demographic, clinical and tissue-related characteristics of the samples. All SZ subjects were institutionalized donors with a long duration of the illness (Table 1, Cohort II) who had no history of neurological episodes. Experienced clinical examiners interviewed each donor ante-mortem to confirm schizophrenia diagnosis according to DSM-IV and International Classification of Diseases 10 (ICD-10) criteria. The last mean daily chlorpromazine equivalent dose for the antipsychotic treatment of patients was based on the electronic records of last drug prescriptions administered up to death (Table 1, Cohort II) and was calculated as previously described [21].

Samples were coded by each brain bank or collection staff to protect human subject confidentiality.

### Protein extraction

Specimens of the lateral cerebellar cortex from the posterior lobe, extending from the pial surface to white matter and only including grey matter, were dissected from coronal slabs stored at −80°C. Protein extracts were prepared from tissue samples using NP40 lysis buffer as described previously [22]. Protein concentration was determined by Bradford assay (Biorad).

### Mass spectrometry screening and data processing

Our screening strategy combines differential isotopic labelling of peptides via reductive dimethylation with offline fractionation by SCX and liquid chromatography coupled to tandem mass spectrometry (LC-MS/MS) on a hybrid linear ion trap mass spectrometer (LTQ-Orbitrap) (Fig S1A). 500 μg of total extracts from four pooled control (100 μg/each) and four pooled schizophrenia (100 μg/each) lysates were digested and further processed as indicated in Supplementary methods. Briefly, the digested peptides were labelled with either hydrogen (light peptides, control) or deuterium (heavy peptides, schizophrenia) isotopes through a reductive dimethylation reaction as described previously [23]. Differentially labelled peptides were then mixed 1:1. Dimethylated peptide mixtures were separated by strong cation exchange (SCX) chromatography on a polysulphoethyl A column. Peptide mixtures were analysed by LC-MS/MS. Each peptide fraction was separated by reverse phase chromatography on a capillary column and analysed online on a hybrid linear ion trap Orbitrap (LTQ-Orbitrap XL, Thermo Scientific) mass spectrometer for identification and relative quantification of pair peptide sequences. MS/MS spectra were searched against a concatenated target-decoy Uniprot human protein database (UP000005640 version 05-23-2017, n=71,567 target sequences) using the Comet search algorithm (version 2015025) and specific search parameters (see Supplementary Methods for more details). The mass spectrometry proteomics data have been deposited in the ProteomeXchange Consortium via the PRIDE partner repository with the dataset identifier PXD008216 [24]. Peptide matches were filtered to <1% False-Discovery Rate (FDR) and protein groups were filtered at ≥90 % probability score. The log_2_ heavy/light ratio for each protein was determined and transformed to a z-score [25]. A significance value (p value) for each protein ratio was calculated from the complementary error function for the normalized distribution of the z-scores [25]. The FDR was computed for all the p values using the Benjamini and Hochberg method [26]. The FDR threshold was set at 0.1 for selected significant proteins with consistent changes amongst peptides. Proteins were classified by their biological function using the Human Protein Reference Database (HPRD-http://www.hprd.org).

The altered observed proteins were compared with those previously reported in proteomic studies of other brain regions of post-mortem samples in schizophrenia based on the gene symbol.

### Immunoblotting

50 μg of total protein lysates were resolved by SDS-PAGE electrophoresis and transferred to a nitrocellulose membrane. Membranes were immunoblotted with rabbit polyclonal antibody against VPP1 (1:500; ab103680, Abcam) and calmodulin (1:1000; 4830, Cell Signalling Technologies), and monoclonal antibodies against parvalbumin (1:1000; MAB1572, Millipore-Chemicon) and glyceraldehyde-3-phosphate dehydrogenase (GAPDH) (1:500000; MAB374, Millipore-Chemicon). All proteins were detected by a unique band at the predicted molecular weight. Densitometric quantification of candidate proteins was performed using Quantity One software (BioRad). Values were normalized to GAPDH and a control reference sample. At least two independent immunoblot analyses were performed per sample.

### Statistical Analysis

For validation analysis using immunoblot we used the following procedures. Normality of the variables was assessed using the Kolmogorov-Smirnov test. Demographic and tissue-related features of the samples were compared between schizophrenia and control conditions using the Fisher exact test for qualitative variables and the Mann-Whitney U test for quantitative non-parametric variables. The differences in protein levels between pools were analysed using the one-tailed unpaired Student’s t test, since the direction of change was expected to be the same as in the proteomics assay. The differences in protein levels between groups in the individual sample analysis were performed using the Kruskal-Wallis one-way analysis of variance by ranks and the Mann-Whitney U test was used to compare differences between two groups. Grubbs test and Pierce test were used to detect outliers for parametrical or non-parametrical variables, respectively; the number of outliers detected for each analysis is indicated in the figure legend. Spearman or Pearson correlation analyses were carried out to detect association of our molecular measures with other clinical, demographic and tissue related variables (age, post-mortem delay, pH and toxicology, and daily antipsychotic dose, age of onset and duration of the illness in the SZ group of Cohort II). Statistical analysis was performed with GraphPad Prism version 5.00, with significance level set to 0.05.

## Results

### Proteomic analysis of post-mortem cerebellum from schizophrenia and control subjects

To identify and quantify disease-related protein changes in schizophrenia, we screened pooled protein lysates from four subjects with schizophrenia and four control subjects matched for gender, age and post-mortem delay (S1A Fig and S1 Table). 1412 proteins with a Protein Prophet probability score of more than 90% were quantified (S1 Data). The distribution of protein heavy-to-light (H/L) ratios revealed that some proteins of the cerebellar proteome were altered in schizophrenia (S1B and S1C Figs). We identified 99 (7%) significant proteins with a false discovery rate acceptance of 10% and a peptide coverage greater than 5% (S2 Data). Moreover, 31 of the 99 significantly regulated proteins were previously reported to be altered in proteomic analyses of other brain areas in schizophrenia (Table 2).

**Table 2.** List of significant altered proteins of the present study that had been previously reported in other brain regions in proteomic studies in schizophrenia.

| **Brain region** | **Gene Symbol** | **Reference\*** |
| --- | --- | --- |
| **DLPFC** | *GNB1, NDUFA12* | (Behan et al., 2009) |
|  | *GNB1, MAG* | (Chan et al., 2011) |
|  | *VIM* | (English et al., 2009) |
|  | *GNB1, TF* | (Martins-de-Souza et al., 2009) |
|  | *ATP6V0D1, SORBS1, VIM* | (Martins-De-Souza et al., 2009) |
|  | *PNP* | (Novikova et al., 2006) |
|  | *TF* | (Pennington et al., 2008) |
|  | *BCL2L13, PHB2, SORBS1, SV2B, TMEM30A* | (Pinacho et al., 2016) |
|  | *TF* | (Prabakaran et al., 2004) |
|  | *PVALB* | (Smalla et al., 2008) |
| **OFC** | *DMTN, PGAM5, VDAC2* | (Velásquez et al., 2017) |
| **ACC** | *GNB1, TF* | (Clark et al., 2006) |
|  | *AP2B1, ATP6V0A1, BAIAP2, CALM2, CCT6A, HSD17B4, NAPA, SRPRB, VDAC2* | (Föcking et al., 2015) |
|  | *GNB1* | (Martins-de-Souza et al., 2010) |
| **CC** | *GNB1, HAPLN2, VIM* | (Saia-Cereda et al., 2015) |
|  | *BAIAP2* | (Saia-Cereda et al., 2016) |
|  | *CALM1, CDC42* | (Saia-Cereda et al., 2017) |
| **Thalamus** | *VIM* | (Martins-de-Souza et al., 2010) |
| **Hippocampus** | *PVALB* | (Föcking et al., 2011) |
|  | *ACOT7, CCT6A* | (Schubert et al., 2015) |
|  | *VIM* | (Nesvaderani et al., 2009) |
| **Temporal lobe** | *HAPLN2* | (Martins-de-Souza et al., 2009) |
|  | *CADM1, CALM1, CDC42, H3F3A, PPP3R1, VIM* | (Saia-Cereda et al., 2017) |
|  | *PHB2, VIM* | (MacDonald et al., 2015) |

DLPFC, dorsolateral prefrontal cortex; OFC, orbitofrontal cortex; ACC, anterior cingulate cortex; CC, corpus callosum. \* The complete information of the references is detailed in the supplementary material.

We further classified the altered proteins according to their biological function and compared them to the non-regulated proteome. Similar biological functions were found in both data sets and transport function was enriched from 10% in the non-regulated proteome to 13% in the regulated proteome (S1D Fig). The most prevalent function was cell communication and signalling pathways (S1D Fig). A list of robust candidates was selected from these functions (Transport and Cell communication and signalling) according to the following criteria: (i) the change had been identified with more than 4 peptides; (ii) a greater than 2-fold increase or decrease. A list of 11 robustly identified altered proteins that belong to these categories was generated (Table 3).

**Table 3.** List of proteins filtered from enriched and most representative functions in the post-mortem cerebellum in schizophrenia.

| Function | Acc. Number | Gene Symbol | Protein Description (Protein Symbol) | Quantified Peptides | Ratio H/L | log<sub>2</sub>Ratio H/L |
| --- | --- | --- | --- | --- | --- | --- |
| **Transport** | **Q93050** | **ATP6V0A1** | **V-type proton ATPase 116 kDa subunit a isoform 1 (VPP1)** | 5 | 0.09 | -3.46 |
|  | A0A0A0MR02 | VDAC2 | Voltage-dependent anion-selective channel protein 2 (VDAC2) | 16 | 0.13 | -2.97 |
|  | Q86UR5 | RIMS1 | Regulating synaptic membrane exocytosis protein 1 (RIMS1) | 11 | 0.46 | -1.12 |
|  | M0R0Y2 | NAPA | Alpha-soluble NSF attachment protein (SNAA) | 6 | 3.90 | 1.97 |
|  | Q01469 | FABP5 | Fatty acid-binding protein, epidermal (FABP5) | 6 | 5.20 | 2.38 |
| **Cell Communication**<br>**/ Signal**<br>**transduction** | **E7EMB3** | **CALM2** | **Calmodulin-2 (CALM2)** | 18 | 2.56 | 1.36 |
|  | P37235 | HPCAL1 | Hippocalcin-like protein 1 (HPCL1) | 9 | 2.64 | 1.40 |
|  | O00533 | CHL1 | Neural cell adhesion molecule L1-like protein (NCHL1) | 17 | 3.01 | 1.59 |
|  | Q8WUD1 | RAB2B | Ras-related protein Rab-2B (RAB2B) | 6 | 3.17 | 1.66 |
|  | Q96FQ6 | S100A16 | Protein S100-A16 (S10AG) | 7 | 3.53 | 1.82 |
|  | **P20472** | **PVALB** | **Parvalbumin alpha (PRVA)** | 8 | 3.65 | 1.87 |

Access numbers from Uniprot database Uniprot; H/L ratio between heavy (schizophrenia) and light (control) peptide areas; selected candidates for validation are shown in bold.

### Validation of hit protein changes in cerebellum schizophrenia samples

Three hit candidates were selected for further validation by immunoblot from the list of robustly determined candidates. From this list, we selected a candidate identified with the highest number of peptides, calmodulin 2 (CALM2), and two more candidates with the most prominent changes in each Gene Ontology function, vacuolar proton translocating ATPase 116 kDa subunit a (VPP1) from transport and parvalbumin alpha (PRVA) from cell communication and signal transduction (Table 3). The antibody for calmodulin detected a pool of calmodulin 1, calmodulin 2 and calmodulin 3, all of which displayed similar molecular weight and protein sequence. We confirmed by immunoblot that the selected candidate changes were also altered in the pooled samples used in proteomics (suicide schizophrenia, n=4; control, n=4, Table S1, S2 Fig). The fold changes of the three candidates were similar to the fold changes detected in the pooled analysis, being decreased for VPP1 and increased for calmodulin and parvalbumin (S2 Fig and Table 3).

We further characterized VPP1, calmodulin and parvalbumin by immunoblot in a cohort consisting of 7 schizophrenia subjects, 7 matched control subjects and 6 matched non-schizophrenia suicide subjects (Table 1, Cohort I). All candidate levels were referred to GAPDH levels, which were not significantly different between the groups (Figs 1A, 1B and 1C). We found that VPP1 protein levels were again significantly decreased [t = 2.809, df = 12, p = 0.0079; Mean ± SEM: control (C) = 1.000 ± 0.143, SZ = 0.526 ± .0.089] and calmodulin and parvalbumin levels were also significantly increased in the schizophrenia group (calmodulin [t = 3.724, df = 12, p = 0.0029; Mean ± SEM: C = 1.000 ± 0.083, SZ = 1.918 ± 0.232], parvalbumin [t = 4.964, df = 12, p = 0.0003; Mean ± SEM: C = 1.000 ± 0.0.086, SZ = 1.524 ± 0.060] (Figs 1A, 1B and 1C). In addition, we found significant differences between the schizophrenia group and the non-schizophrenia suicide group in the fold changes for calmodulin and parvalbumin, which in the suicide group showed similar levels to the control group [calmodulin [t = 2.708, df = 11, p = 0.0204; Mean ± SEM: SZ = 1.918 ± 0.232, suicide (SC) = 1.141 ± 0.149], parvalbumin [t = 6.129, df = 11, p < 0.0001; Mean ± SEM: SZ = 1.524 ± 0.060, SC = 0.849 ± 0.096] (Figs 1A, 1B and 1C). However, the protein levels of VPP1 were not significantly different in the non-schizophrenia suicide group compared to the schizophrenia group [t = 0.3354, df = 11, p = 0.7436; Mean ± SEM: SZ = 0.526 ± 0.089, SC = 0.492 ± 0.032] (Figs 1A, 1B and 1C). Furthermore, we analysed the influence of other demographic, clinical and tissue-related variables in the differences found in the schizophrenia group compared to control subjects. None of our molecular measures showed any association with other variables of the study with the exception of parvalbumin, which showed an increase in the antipsychotic treated group compared to drug-free subjects (Table 4, Cohort I; Dunn’s Test p<0.05), indicating that the increase observed for parvalbumin in schizophrenia could be due to the antipsychotic treatments in these subjects.

**Table 4:** Association analysis of other variables in Cohorts I and II.

| Cohort I |  |  |  |  |
| --- | --- | --- | --- | --- |
|  | Age | PMD | pH | Toxicology |
|  | r | r | r | K |
| C-SZ cohort (n = 14) |  |  |  |  |
| VPP1 | 0.095 | -0.121 | -0.226 | 5.202 |
| CaM<sup>#</sup> | -0.035 | -0.412 | 0.239 | 0.733 |
| PRVA | 0.107 | 0.015 | 0.071 | **6.959<sup>a</sup>** |
| Cohort II |  |  |  |  |
|  | Age | PMD | pH |  |
|  | r | r | r |  |
| SZ-C cohort II (n=27) |  |  |  |  |
| CaM<sup>†</sup> | 0.221 | -0.171 | -0.048 |  |
| PRVA | 0.032 | 0.208 | 0.169 |  |
|  | Daily AP dose<sup>‡</sup> | Age of onset | Duration of illness |  |
|  | r | r | r' |  |
| SZ cohort II (n=13) |  |  |  |  |
| CaM | -0.116 | -0.099 | 0.124 |  |
| PRVA | 0.162 | -0.170 | -0.028 |  |

r, Pearson's r for parametric variables; r', Spearman's correlation for non-parametric variables; PMD, post-mortem delay; C, control; SZ, schizophrenia; AP, antipsychotic. #r of Spearman are shown for this protein. <sup>†</sup> An outlier was detected for calmodulin (CaM) and therefore excluded from the SZ-C analysis (CaM: C, n=13, SZ, n=13). <sup>‡</sup>Last chlorpromazine equivalent dose was calculated based on the electronic records of drug prescriptions of the patients; K, Kruskal-Wallis; N/A, not applicable. Significant associations are indicated in bold.
<sup>a</sup>p<0.05.

**Figure 1.**
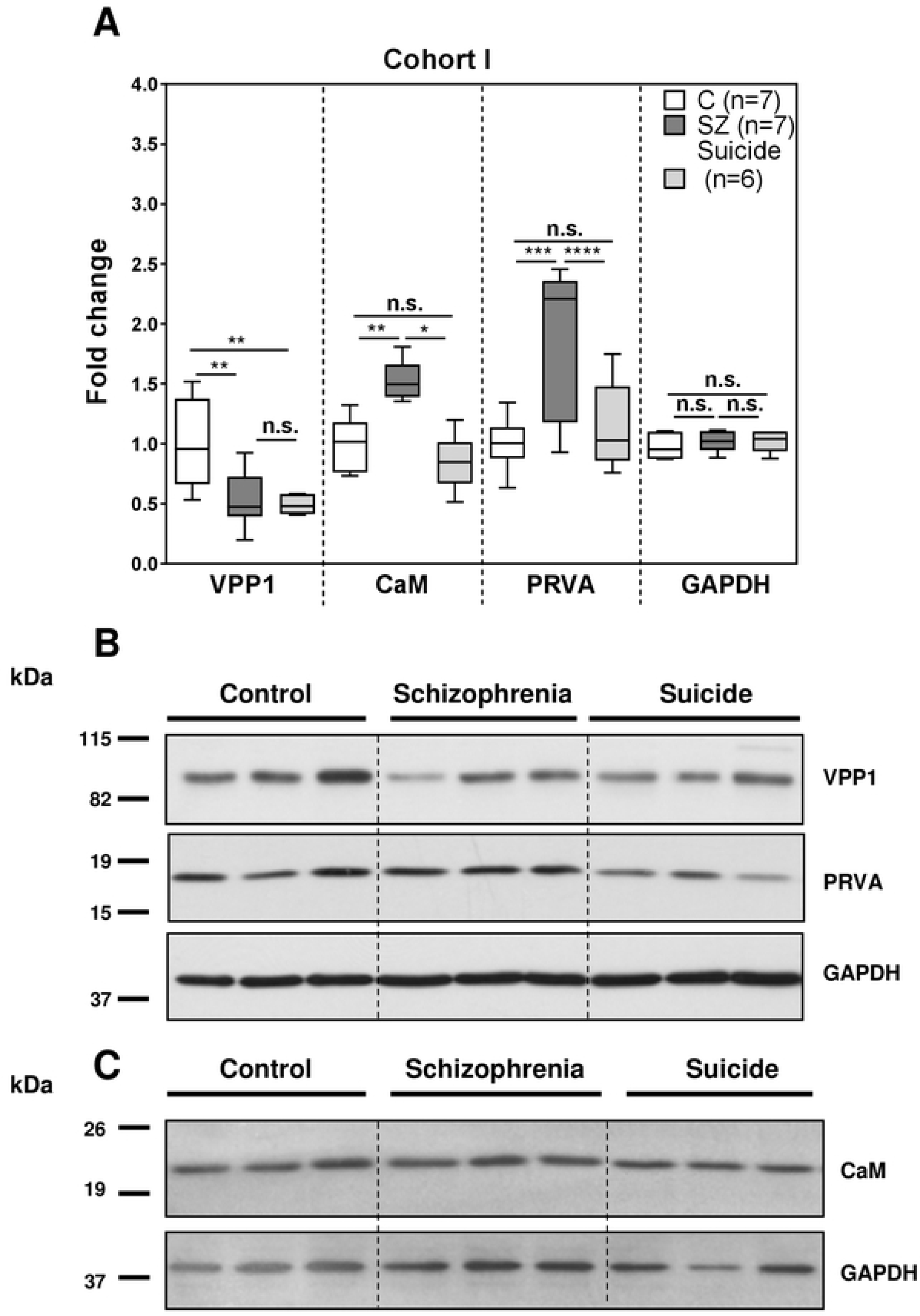
Validation analysis of hit candidate proteins by immunoblot in cohort I. Protein extracts from samples of the post-mortem cerebellum of non-psychiatric control (C, n=7), schizophrenia (SZ, n=7) and non-schizophrenia suicide (n=6) subjects (Table 1, Cohort I) were analysed by immunoblot for VPP1, PRVA, calmodulin (CaM) and GAPDH and quantified by densitometry. **(A)** Protein levels for each were normalized to GAPDH values and to the mean of the control samples. Each box plot represents the median, interquartile range and range of each group. Statistical analysis was performed using the Kruskal-Wallis test for differences between groups (VPP1: p=0.0095; PRVA: p=0.0011; CaM: p=0.0270) and the t test for comparison between the indicated groups. (n.s.-not significant, *p<0.05, **p<0.01, ***p<0.001, ****p<0.0001). **(B)** Representative Western blot images for VPP1, PRVA and GAPDH in 3 non-psychiatric control individuals, 3 schizophrenia subjects and 3 non-schizophrenia suicide controls. **(C)** Representative Western blot images for CaM and GAPDH in 3 non-psychiatric control individuals, 3 schizophrenia subjects and 3 non-schizophrenia suicide control. See supplementary Figure S3 for complete western blot images.

We further characterized VPP1, calmodulin and parvalbumin protein levels by immunoblot in an independent and larger cohort of 13 non-suicide chronic schizophrenia subjects and 14 matched control individuals (Table 1, Cohort II). All candidate levels were referred to GAPDH levels, which were not significantly different between the groups (Figs 2A, 2B and 2C). We found that calmodulin and parvalbumin protein levels were significantly altered in the non-suicide schizophrenia group (calmodulin [U = 45.00, p = 0.0136; Mean ± SEM: C = 1.000 ± 0.3570, SZ = 0.6586 ± 0.2235], parvalbumin [t = 2.337, df = 25, p = 0.0139; Mean ± SEM: C = 1.000 ± 0.110, SZ = 1.380 ± 0.163]) (Figs 2A, 2B and 2C). VPP1 protein levels did not show a significant decrease in the schizophrenia group compared to control group [t = 0.1384, df = 24, p = 0.4455; Median ± SEM: C = 0.844 ± 0.148, SZ = 0.884 ± 0.249] (Figs 2A, 2B and 2C). Association analysis of other variables in this cohort of chronic schizophrenia (age, post-mortem *delay*, pH, antipsychotic dose, age of onset and duration of the illeness) did not show any associations between calmodulin and parvalbumin protein levels and other variables (Table 4, Cohort II).

**Figure 2.**
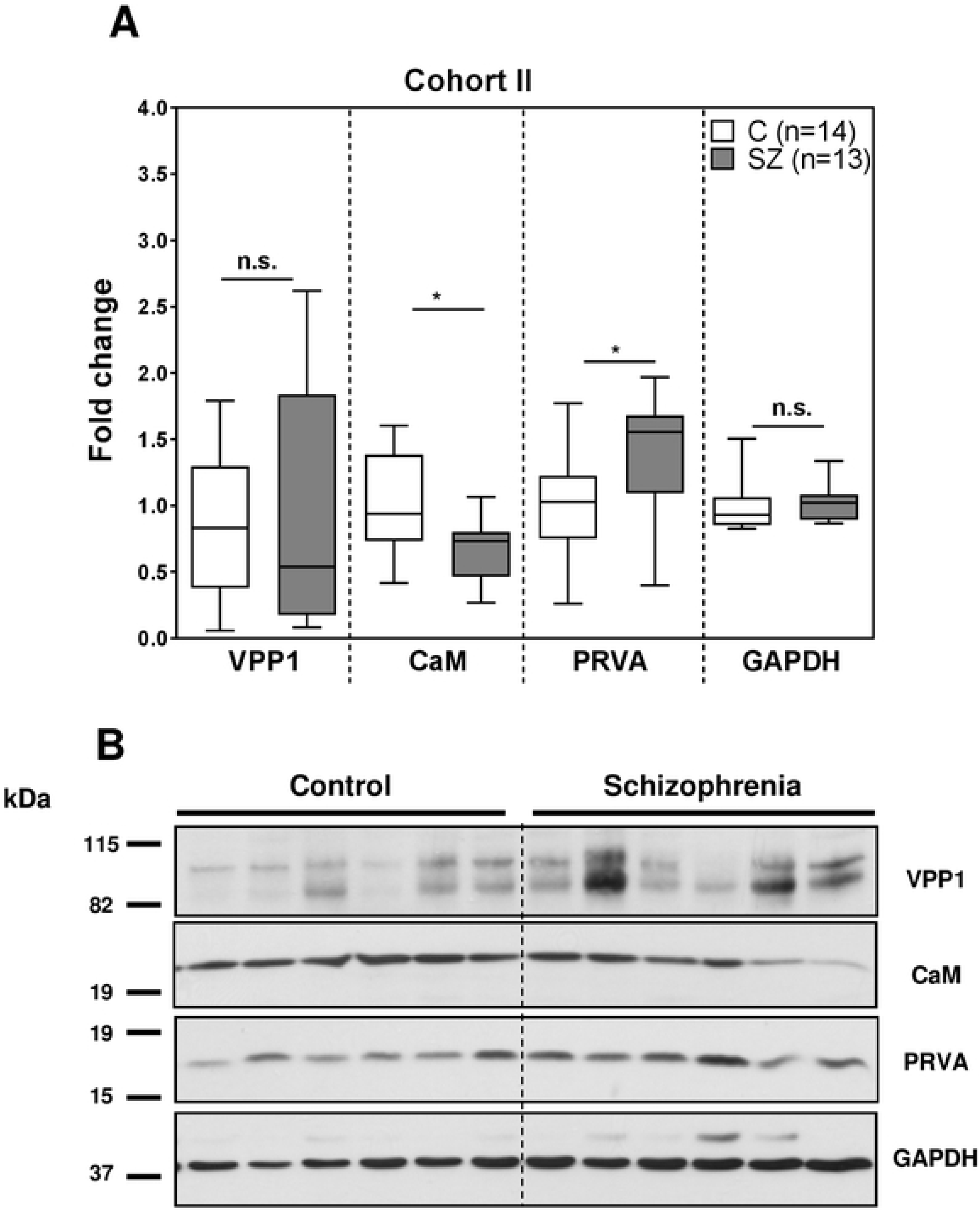
Validation analysis of hit candidate protein by immunoblot in cohort II. Protein extracts from samples of the post-mortem cerebellum of non-suicide chronic schizophrenia (SZ, n=13) and control (C, n=14) subjects (Table 1, Cohort II) were analysed by immunoblot for the same proteins as in Fig 1A and quantified by densitometry. **(A)** Protein levels for each protein were normalized to GAPDH values and to the mean of control samples. Each box plot represents the median, interquartile range and range of each group. An outlier was detected for VPP1 proteins values in the control group. Statistical analysis for comparison between case and control groups was performed using the t test for VPP1 and PRVA and the Mann-Whitney U test for CaM and GAPDH (n.s.-not significant, *p<0.05, ***p<0.001). **(B)** Representative Western blot images for VPP1, PRVA, CaM and GAPDH in 6 non-psychiatric control individuals and 6 non-suicide schizophrenia subjects from the set 2. See supplementary Figure S4 for complete western blot images.

## Discussion

Proteomic screening of cerebellum samples from schizophrenia patients has helped to identify candidates with a putative relevant role in this disorder. The list of altered proteins in this region in suicide schizophrenia subjects provides the first step to analyse in depth some of the most robust candidates. A systematic validation of three candidates in two independent cohorts, one containing suicide individuals and another containing schizophrenia subjects that died of natural causes, revealed that only the two EF-hand calcium-binding proteins were altered in schizophrenia but not in non-schizophrenia suicide subjects, while the third candidate, a subunit of the proton pump ATPase, was linked to suicide behaviours and was not dysregulated in non-suicidal schizophrenia subjects. Thus, our findings propose that the EF-hand calcium-binding proteins parvalbumin and calmodulin could be involved in the pathophysiology of schizophrenia in the cerebellum by altering calcium-dependent signalling pathways involved in synaptic function. Moreover, this work provides evidence for the role of VPP1 in suicide behaviours with a possible impact on synaptic vesicle cycle. Both alterations in the calcium-binding proteins in schizophrenia and in the proton pump ATPase in suicide behaviours could lead to a disruption in the cerebellar synaptic functioning of the neuronal circuits through different mechanisms as discussed below.

### Vacuolar proton ATPase pump

Here we describe a decrease in protein levels of the subunit of the vacuolar-type proton pump ATPase (VPP1). VPP1 in suicide subjects with schizophrenia or other psychiatric disorders, but not in non-suicidal elderly schizophrenia patients, suggesting that this protein may be involved in suicide behaviours rather than in schizophrenia. Another report has also found a decrease in VPP1 in younger schizophrenia subjects in the anterior cingulated cortex using the Stanley Medical Research Institute’s (SMRI) Array Collection (http://www.stanleyresearch.org), which included 4 suicides out of 15 subjects [27]. In line with our results, VPP1 has been reported in a list of genes differently expressed in the frontal cortex in subjects with major depression that committed suicide [28]. VPP1 is part of the large multi-subunit complex of the vacuolar proton pump ATPase. This complex provides a proton motive force and is involved in many cellular and physiological functions (reviewed in [29]). V^+^H-ATPase pumps are composed of two functional components, the cytoplasmic V1 component with the catalytic activity for ATP hydrolysis and the V0 component, which forms a membrane embedded component that is required for proton translocation across the membrane, which occurs through the a subunit (reviewed in [29]). The VPP1subunit is highly expressed in neuronal cells. This subunit is a major component of synaptic vesicles and provides the pH gradient and membrane potential required for neurotransmitter accumulation in the initial phase of the synaptic vesicle cycle [30,31]. Later studies provided evidence that subunit a of the V0 domain is also involved in synaptic vesicle release by mediating the membrane fusion events downstream of the t-SNARE docking of vesicles in a calcium/calmodulin-dependent manner [32,33]. Indeed, the VPP1 orthologue in fly neurons directly interacts with calmodulin, another protein that we found altered in schizophrenia (see above), and this interaction is required for recruiting calmodulin to synapses and for the viability function of VPP1 [32,34]. In addition, neurons lacking VPP1 in *Drosophila melanogaster* and *Caenorhabditis elegans* accumulate vesicles in the synaptic terminals, supporting the role of this protein in neurotransmitter release into the synaptic cleft [32,35,36]. Thus, the reduction of VPP1 that we found in the cerebellum in suicide subjects could be impairing normal neurotransmitter accumulation in synaptic vesicles and their subsequent release into the synaptic terminals. A limited synaptic response of the cerebellar neurons could contribute to the altered functioning of cerebellar circuits in suicide behaviours.

In agreement with this idea of altered synaptic activity in suicide are the findings of multiple studies. The regulation of the expression of genes involved in neurotransmission and synaptic function has been linked to suicide behaviours in different psychiatric disorders including schizophrenia and major depression [28,37– 40]. Interestingly, the cerebellum has been described as one of the neural routes altered in suicide behaviours related to monoaminergic signal transduction [41]. Some reports have also linked reduced cerebellar activity and reduced grey matter with the ideation of suicide or suicide attempt, respectively [42,43]. Thus, the reduction of VPP1 observed in our study in suicide subjects in the cerebellum could alter neurotransmitter refilling of vesicles and their release in cerebellar circuits. Moreover, it might mediate the decreased activity reported in this area in suicide behaviours.

### EF-hand calcium-binding proteins

Calcium homeostasis has been suggested to be disrupted in schizophrenia [44,45]. In our study, we found altered levels of two proteins that are sensitive to changes in the intracellular concentrations of calcium and that play a key role in signalling transduction and synaptic functioning.

#### Calmodulin

Calmodulin (CaM) belongs to the large family of EF-hand calcium-binding proteins. Here, we report an increase in calmodulin levels in the cerebellum in schizophrenia in chronic SZ group with a mean age of 44 years while a downregulation for calmodulin was found in an elderly chronic schizophrenia cohort (mean age of 74 years). In a previous study in elderly chronic schizophrenia has also been reported a downregulation of calmodulin protein levels in different brain areas [46]. In a study of chronic schizophrenia, an upregulation of protein levels of calmodulin has also been reported in cohort with a mean age around 60 years in corpus callosum and anterior temporal lobe [47]. In line with our study, these reports suggest a different regulation of calmodulin depending on the age. Calmodulin is the major calcium-binding protein present in the brain and acts as a calcium sensor, detecting and responding to biologically relevant changes in intracellular concentrations of calcium [48–50]. Calcium regulates calmodulin by changing its subcellular localization, promoting interaction with many proteins or by inducing conformational changes that allow the interaction and activation of specific targets and the subsequent triggering of a signalling cascade [48]. Calmodulin-dependent kinase II (CaMKII) has been proposed as a susceptibility gene in schizophrenia [51]. CaMKII is an important calmodulin effector in neurons that has been implicated in activity-dependent functions such as gene transcription, signalling and synaptic and dendritic development, maturation and function as well as in cognition [49,52]. The list of calmodulin effectors is extensive and includes plasma membrane calcium pumps, various ion channels, protein kinases and receptors, among others [48,50,53]. Thus, the calmodulin-dependent functions that could be dysregulated in schizophrenia in the cerebellum are wide, leading to an important impact on cerebellar circuit functioning and formation in accordance with the connectivity deficits and the neurodevelopmental hypothesis for schizophrenia [1,54]. In our study, we also found in suicide schizophrenia subjects a decrease in one interactor of calmodulin in flies, the orthologue of VPP1, which is required for recruiting calmodulin to synapses [34,55]. Indeed, the Ca^2+^-CaM regulation of V100 (VPP1 orthologue in flies) has been proposed as a positive regulator of the assembly of the SNARE complex on distinct vesicles and subsequent neurotransmitter release [34]. If this interaction is present in cerebellar neurons, this evidence could either suggest that the upregulation of calmodulin in the cerebellum in suicide schizophrenia subjects could be a mechanism to compensate for the possible lower abundance of calmodulin at the synapses due to a reduction in VPP1 levels or that it could be a compensatory mechanism for the lower formation/release of synaptic vesicles [34,55]. Further investigations will be needed to confirm this possibility. However, in SZ patients that die by natural causes, calmodulin was decrease, suggesting a possible constitutive downregulation of calmodulin in schizophrenia. In this context, VPP1 levels were not altered, raising the possibility that the recruitment of calmodulin to the synapse could be correct and the reduction of calmodulin could be impacting in other calmodulin-dependent functions in the cerebellum.

#### Parvalbumin

Here, we report an increase in parvalbumin in the cerebellum in schizophrenia independently of the mechanism of death. Parvalbumin, like calmodulin, also belongs to the large family of EF-hand calcium-binding proteins. However, parvalbumin is a calcium buffer protein, which are proteins essential for modulating calcium homeostasis in neurons and are implicated in the subtle regulation and timing of calcium signals pre- or postsynaptically [56–58]. Parvalbumin is a spatial and temporal regulator of calcium transients that modulates calcium pools and that is critical for synaptic plasticity, such as short-term facilitation [56,57]. Furthermore, it has been proposed that it could also regulate calcium signalling as a slow-onset calcium sensor in addition to being regulated by calcium [59]. Parvalbumin is expressed in subpopulations of GABAergic interneurons in the brain, which are considered more metabolically and electrically active that the neighbouring neurons and which play an important role in the pathophysiology of schizophrenia [12,60,61]. Indeed, a decrease in the expression of parvalbumin in patients with SZ in different brain regions including the hippocampus and prefrontal cortex has been a consistent finding in human post-mortem studies [45,62] and in animal models of schizophrenia [63–66]. However, little is known about parvalbumin protein levels in the cerebellum of SZ patients. In the cerebellum, parvalbumin locates in the axon, soma and dendrites of Purkinje, stellate, basket and a small proportion of Golgi cells [59]. Studies in parvalbumin-deficient mice suggest that this protein is required for normal locomotor activity in order to maintain the normal spontaneous arrhythmic and asynchronous firing pattern of Purkinje cells [67,68]. These mice also showed behavioural deficits linked to schizophrenia and autism, such as deficits in sensorimotor gating and novelty seeking and reduced social interaction and communication [69,70]. The increase in parvalbumin observed in this study in the cerebellum in schizophrenia could produce an imbalance in the regulation of intracellular calcium concentration and/or altering calcium-dependent signalling during synapse activity and this could have an impact on the behavioural changes observed in schizophrenia. The increase in parvalbumin could also reflect a change in the number of certain subpopulations of parvalbumin-positive neurons in the cerebellum in schizophrenia. However, a reduction in Purkinje neurons in the cerebellum has been reported in subjects with schizophrenia [61], suggesting that this possibility may be more likely due to other types of parvalbumin-expressing neurons in the cerebellum. Further studies will be needed to elucidate these possibilities.

Functional neuroimaging studies show a predominantly hypoactivation of cerebellum in the schizophrenia [71,72]. Thus, based on our results, the altered EF-hand calcium-binding proteins found in the cerebellum could have an impact on synaptic transmission and underlie the reduced cerebellar activity observed in people with schizophrenia. Further studies are needed to investigate this hypothesis.

### Limitations

We acknowledge several limitations of our study. Firstly, we used pooled samples in the proteomic screening. Although this type of design is a useful approach for detecting common altered pathways [73–77], it does not allow for the control of inter-individual variations, which could account for modifications in the resulting molecular signature. However, in our analysis the results obtained by immunoblot in pools were also observed in the individual sample analysis in a similar cohort, suggesting that molecular changes in the cerebellum could be stable between individuals. Secondly, the sample size of Cohort I is limited. Further analysis in an independent and larger cohort of samples including suicidal groups will be needed to explore how stable our findings are in other patients. Thirdly, antipsychotic treatments could influence the results. To control for this variable, we have used the blood toxicology data in Cohort I and chlorpromazine daily dose data in Cohort II, showing that the increased expression of parvalbumin could be due to the antipsychotic treatments in Cohort I but not in Cohort II. Further pharmacological studies in cellular and animal models, as well as in drug-naive patients, will help to clarify the effect of antipsychotics on parvalbumin. Fourthly, the schizophrenia subjects used in our proteomic screening committed suicide, which could be influencing the findings observed. We have controlled for this possibility in the validation of candidates by including a group that includes subjects who committed suicide but with varying psychiatric diagnoses instead of schizophrenia and an independent cohort with non-suicide schizophrenia subjects that died by natural causes. These analyses have allowed us to detect that VPP1 alteration could be a common feature of suicide and suggest that this candidate could be only altered in suicide subjects rather than in schizophrenia. It could also be possible that alterations of these proteins could occur in other psychiatric disorders. Further studies of these candidates in anxiety and adjustment disorders will be helpful to explore this possibility.

## Conclusions

In summary, our findings provide evidence for an upregulation of calcium-binding proteins in the cerebellum in schizophrenia, suggesting an altered modulation of calcium signalling and calcium transients in synapse response in this region in schizophrenia with an impact on the normal functioning of cerebellar circuits. In addition, our study provides evidence for the alteration of VPP1 in the cerebellum linked to suicide behaviours, suggesting an involvement of defective synaptic vesicle cycle and the release of neurotransmitters in suicide behaviours.

## Abbreviations

SZ: Schizophrenia
SCX: Strong cation exchange
LTQ: Linear Trap Quadrupole (mass spectrometer)
DSM-IV: Diagnostic and Statistical Manual of Mental Disorders-Fourth Edition
PMD: Post-mortem delay
ICD-10: International Classification of Diseases 10
ACN: Acetonitrile
VPP1: Vacuolar proton pump 1
PRVA: Parvalbumin alpha
CALM2: Calmodulin 2
CaM: Calmodulin

## Acknowledgements

These studies were supported by funding from a Miguel Servet grant (MS16/00153-CP16/00153) to BR financed and integrated into the National R + D + I and funded by the ISCIII - General Branch Evaluation and Promotion of Health Research - and the European Regional Development Fund (ERDF), the Basque Government (IT1211/19) and SAF 2017-88126-R to J.J.M.

The authors thank the donors and their families for the donation of their brains; the technical support of Nuria Villalmanzo; the staff members of the Basque Institute of Legal Medicine for their help; and the English language correction of Dr. Rose.

## Supporting Information Legends

**Figure S1. Experimental strategy for large-scale quantitative proteomic analysis and identification of differentially expressed proteins in cerebellum in schizophrenia. (A)** Protein lysates from the post-mortem cerebellum of control (n=4) and suicide schizophrenia (SZ, n=4) subjects (Table 1) were processed as described in the experimental procedures section. In the analysis in pools, samples from the same group were pooled. In the analysis of individual samples from schizophrenia, a pool of controls was used to compare each individual sample of schizophrenia. Subsequently, protein database searches, peptide quantification and data analysis were performed as described in the experimental procedures section. A panel of 3 candidates from significantly regulated proteins was selected for further validation by immunoblot: first, in a pilot cohort which includes a group of non-schizophrenia suicide subjects (Table 1; Cohort I: SZ (n =7), non-SZ suicide (n=6), control (n=7)) and then, in a larger cohort of non-suicide chronic schizophrenia subjects (Table 1, Cohort II: non-suicide SZ (n=13), control (n=14)). **(B)** Distribution of the number of peptides quantified per protein from the data set of 2289 quantified proteins. **(C)** Normalized distribution of z-scores for confidently quantified proteins (>2 peptide sequences) (n=1148). **(D)** Gene ontology classification of biological functions for non-significantly and significantly altered proteins with low variation in cerebellum in SZ compared to control. Transport (GO:0006810); Cell communication (GO:0007154); Signal transduction (GO:0007165); Metabolism (GO:0008152); Energy pathways (GO:0006091); Regulation of nucleobase, nucleoside, nucleotide and nucleic acid metabolism (GO:0019219); Cell growth and/or maintenance (GO:0008151); Protein metabolism (GO:0019538); Biological process unknown (GO:0000004).

**Figure S2. Validation of hit candidate proteins by immunoblot in pools.** Pooled protein extracts from samples of the post-mortem cerebellum of control (C, n=4), and schizophrenia (SZ, n=4) subjects from the *UPV/EHU brain collection* (Table S1, a subgroup from Cohort I, Table 1) used in the proteomic screening were analysed by immunoblotting for ATP6V0A1, PVALB, calmodulin (CaM) and GAPDH. Protein levels for each hit were quantified by densitometry and normalized to GAPDH values and to the reference control sample. Images show representative immunoblots of a pool of control (left band, C) and a pool of schizophrenia (right band, SZ) subjects. Analysis was performed in duplicate. Bars represent mean ± standard deviation of the analysis of duplicates from two independent dissections, with the exception of PVALB, whose data are from a duplicate analysis of one dissection. Statistical analysis was performed using t test (n.s.-not significant, **p<0.01, ***p<0.001).

**Figure S3. Images of completed Westerns blot membranes from Figure 1.** Completed images of Western blot showed in Figure 1. Completed images of anti-VVP1 (A), anti-GAPDH (B), anti-PRVA (C) or anti-calmodulin (D) immunoreactivity. The cut of the membranes of western blot are shown by round dot lines. The samples shown in Figure 1 are limited by dashed line at the complete membranes of western blot. *, Immunoreactivity of other antibodies.

**Figure S4. Images of completed Westerns blot membranes from Figure 2.** Completed images of Western blot showed in Figure 1. Completed images of anti-VVP1 and anti-GAPDH (A), anti-PRVA (B) or anti-calmodulin (C) immunoreactivity. The cut of the membranes of western blot are shown by round dot lines. The samples shown in Figure 2 are limited by dashed line at the complete membranes of western blot. *, Immunoreactivity of other antibodies.

## References

1. Andreasen NC, Nopoulos P, O’Leary DS, Miller DD, Wassink T, Flaum M. Defining the phenotype of schizophrenia: cognitive dysmetria and its neural mechanisms. Biol Psychiatry. 1999;46: 908–20. Available: doi: 10.1016/s0006-3223(99)00152-3

2. Parker KL, Kim YC, Kelley RM, Nessler AJ, Chen K-H, Muller-Ewald VA, et al. Delta-frequency stimulation of cerebellar projections can compensate for schizophrenia-related medial frontal dysfunction. Mol Psychiatry. 2017;22: 647–655. Available: doi: 10.1038/mp.2017.50

3. Brady RO, Gonsalvez I, Lee I, Öngür D, Seidman LJ, Schmahmann JD, et al. Cerebellar-prefrontal network connectivity and negative symptoms in schizophrenia. Am J Psychiatry. 2019. Available: doi: 10.1176/appi.ajp.2018.18040429

4. Adamaszek M, D’Agata F, Ferrucci R, Habas C, Keulen S, Kirkby KC, et al. Consensus paper: cerebellum and emotion. The Cerebellum. 2017;16: 552–576. Available: doi: 10.1007/s12311-016-0815-8

5. Andreasen NC, Pierson R. The Role of the cerebellum in schizophrenia. Biol Psychiatry. 2008;64: 81–88. Available: doi: 10.1590/S1807-59322011001300009

6. Sokolov AA, Miall RC, Ivry RB. The cerebellum: Adaptive prediction for movement and cognition. Trends Cogn Sci. 2017;21: 313–332. Available: doi: 10.1016/j.tics.2017.02.005

7. Mothersill O, Knee-Zaska C, Donohoe G. Emotion and theory of mind in schizophrenia—investigating the role of the cerebellum. The Cerebellum. 2016;15: 357–368. Available: doi: 10.1007/s12311-015-0696-2

8. Ben-Yehudah G, Guediche S, Fiez JA. Cerebellar contributions to verbal working memory: beyond cognitive theory. The Cerebellum. 2007;61. Ben-Ye: 193–201. Available: doi: 10.1080/14734220701286195

9. Bellebaum C, Daum I. Cerebellar involvement in executive control. The Cerebellum. 2007;6: 184–192. Available: doi: 10.1080/14734220601169707

10. Ackermann H, Mathiak K, Riecker A. The contribution of the cerebellum to speech production and speech perception: Clinical and functional imaging data. The Cerebellum. 2007;6: 202–213. Available: doi: 10.1080/14734220701266742

11. Fatemi SH, Folsom TD. GABA receptor subunit distribution and FMRP–mGluR5 signaling abnormalities in the cerebellum of subjects with schizophrenia, mood disorders, and autism. Schizophr Res. 2015;167: 42–56. Available: doi: 10.1016/j.schres.2014.10.010

12. Fatemi SH, Folsom TD, Rooney RJ, Thuras PD. Expression of GABAA α2-, β1- and ɛ-receptors are altered significantly in the lateral cerebellum of subjects with schizophrenia, major depression and bipolar disorder. Transl Psychiatry. 2013;3: e303–e303. Available: doi: 10.1038/tp.2013.64

13. Burnet P, Hutchison L, Vonhesling M, Gilbert E, Brandon N, Rutter A, et al. Expression of D-serine and glycine transporters in the prefrontal cortex and cerebellum in schizophrenia. Schizophr Res. 2008;102: 283–294. Available: doi: 10.1016/j.schres.2008.02.009

14. Schmitt A, Koschel J, Zink M, Bauer M, Sommer C, Frank J, et al. Gene expression of NMDA receptor subunits in the cerebellum of elderly patients with schizophrenia. Eur Arch Psychiatry Clin Neurosci. 2010;260: 101–111. Available: doi: 10.1007/s00406-009-0017-1

15. Pinacho R, Villalmanzo N, Roca M, Iniesta R, Monje A, Haro JM, et al. Analysis of Sp transcription factors in the postmortem brain of chronic schizophrenia: A pilot study of relationship to negative symptoms. J Psychiatr Res. 2013;47: 926–934. Available: doi: 10.1016/j.jpsychires.2013.03.004

16. Keller S, Punzo D, Cuomo M, Affinito O, Coretti L, Sacchi S, et al. DNA methylation landscape of the genes regulating D-serine and D-aspartate metabolism in post-mortem brain from controls and subjects with schizophrenia. Sci Rep. 2018;8: 10163. Available: doi: 10.1038/s41598-018-28332-x

17. Mudge J, Miller NA, Khrebtukova I, Lindquist IE, May GD, Huntley JJ, et al. Genomic convergence analysis of schizophrenia: mRNA sequencing reveals altered synaptic vesicular transport in post-mortem cerebellum. PLoS One. 2008;3: e3625. Available: doi: 10.1371/journal.pone.0003625

18. Kim S, Cho H, Lee D, Webster MJ. Association between SNPs and gene expression in multiple regions of the human brain. Transl Psychiatry. 2012;2: e113. Available: doi: 10.1038/tp.2012.42

19. Kim S, Webster MJ. The Stanley Neuropathology Consortium Integrative Database: a novel, web-Based tool for exploring neuropathological markers in psychiatric disorders and the biological processes Associated with Abnormalities of Those Markers. Neuropsychopharmacology. 2010;35: 473–482. Available: doi: 10.1038/npp.2009.151

20. Roca Casasús M, Escanilla Casal A, Monje Hernández A, Baño Galindo V, Planchat Teruel LM, Costa Escola J, et al. Banco de tejidos neurológicos de Sant Joan de Déu-Serveis de Salut Mental para la investigación de las enfermedades mentales. La importancia de un programa de donación en vida. Psiquiatr Biológica. 2008;15: 73–79. Available: https://doi.org/10.1016/S1134-5934(08)71126-6

21. Gardner DM, Murphy AL, O’Donnell H, Centorrino F, Baldessarini RJ. International consensus study of antipsychotic dosing. Am J Psychiatry. 2010;167: 686–693. Available: doi: 10.1176/appi.ajp.2009.09060802

22. Pinacho R, Villalmanzo N, Lalonde J, Haro JM, Meana JJ, Gill G, et al. The transcription factor SP4 is reduced in postmortem cerebellum of bipolar disorder subjects: control by depolarization and lithium. Bipolar Disord. 2011;13: 474–485. Available: doi: 10.1111/j.1399-5618.2011.00941.x

23. Khidekel N, Ficarro SB, Clark PM, Bryan MC, Swaney DL, Rexach JE, et al. Probing the dynamics of O-GlcNAc glycosylation in the brain using quantitative proteomics. Nat Chem Biol. 2007;3: 339–348. Available: doi: 10.1038/nchembio881

24. Vizcaíno JA, Csordas A, Del-Toro N, Dianes JA, Griss J, Lavidas I, et al. 2016 update of the PRIDE database and its related tools. Nucleic Acids Res. 2016;44: D447–D456. Available: doi: 10.1093/nar/gkv1145

25. Graumann J, Hubner NC, Kim JB, Ko K, Moser M, Kumar C, et al. Stable isotope labeling by amino acids in cell culture (SILAC) and proteome quantitation of mouse embryonic stem cells to a depth of 5,111 proteins. Mol Cell Proteomics. 2008;7: 672–683. Available: doi: 10.1074/mcp.M700460-MCP200

26. Benjamini Y, Hochberg Y. Controlling the false discovery rate: a practical and powerful approach to multiple testing. J Roy Stat Soc Ser B. 1995. pp. 289–300. Available: https://doi.org/10.1111/j.2517-6161.1995.tb02031.x

27. Föcking M, Lopez LM, English JA, Dicker P, Wolff A, Brindley E, et al. Proteomic and genomic evidence implicates the postsynaptic density in schizophrenia. Mol Psychiatry. 2015;20: 424–432. Available: doi: 10.1038/mp.2014.63

28. Zhurov V, Stead JDH, Merali Z, Palkovits M, Faludi G, Schild-Poulter C, et al. Molecular pathway reconstruction and analysis of disturbed gene expression in depressed individuals who died by suicide. Yaragudri VK, editor. PLoS One. 2012;7: e47581. Available: doi: 10.1371/journal.pone.0047581

29. Maxson ME, Grinstein S. The vacuolar-type H+-ATPase at a glance - more than a proton pump. J Cell Sci. 2014;127: 4987–4993. Available: doi: 10.1242/jcs.158550

30. Moriyama Y, Futai M. H(+)-ATPase, a primary pump for accumulation of neurotransmitters, is a major constituent of brain synaptic vesicles. Biochem Biophys Res Commun. 1990;173: 443–8. Available: doi: 10.1016/s0006-291x(05)81078-2

31. Takamori S, Holt M, Stenius K, Lemke EA, Grønborg M, Riedel D, et al. Molecular Anatomy of a Trafficking Organelle. Cell. 2006;127: 831–846. Available: doi: 10.1016/j.cell.2006.10.030

32. Morel N, Poëa-Guyon S. The membrane domain of vacuolar H+ATPase: a crucial player in neurotransmitter exocytotic release. Cell Mol Life Sci. 2015;72: 2561–2573. Available: doi: 10.1007/s00018-015-1886-2

33. Bayer MJ, Reese C, Bühler S, Peters C, Mayer A. Vacuole membrane fusion: V0 functions after trans-SNARE pairing and is coupled to the Ca2+-releasing channel. J Cell Biol. 2003;162: 211–222. Available: doi: 10.1083/jcb.200212004

34. Wang D, Epstein D, Khalaf O, Srinivasan S, Williamson WR, Fayyazuddin A, et al. Ca2+ –Calmodulin regulates SNARE assembly and spontaneous neurotransmitter release via v-ATPase subunit V0a1. J Cell Biol. 2014;205: 21–31. Available: doi: 10.1083/jcb.201312109

35. Hiesinger PR, Fayyazuddin A, Mehta SQ, Rosenmund T, Schulze KL, Zhai RG, et al. The v-ATPase V0 subunit a1 is required for a late step in synaptic vesicle exocytosis in Drosophila. Cell. 2005;121: 607–620. Available: doi: 10.1016/j.cell.2005.03.012

36. Liégeois S, Benedetto A, Garnier JM, Schwab Y, Labouesse M. The V0-ATPase mediates apical secretion of exosomes containing Hedgehog-related proteins in Caenorhabditis elegans. J Cell Biol. 2006;173: 949–961. Available: doi: 10.1083/jcb.200511072

37. Roy B, Dwivedi Y. Understanding the neuroepigenetic constituents of suicide brain. Progress in molecular biology and translational science. 2018. pp. 233–262. doi:10.1016/bs.pmbts.2018.01.007

38. Serafini G, Pompili M, Innamorati M, Giordano G, Montebovi F, Sher L, et al. The role of microRNAs in synaptic plasticity, major affective disorders and suicidal behavior. Neurosci Res. 2012;73: 179–190. Available: doi: 10.1016/j.neures.2012.04.001.

39. Bortolato M, Pivac N, Muck Seler D, Nikolac Perkovic M, Pessia M, Di Giovanni G. The role of the serotonergic system at the interface of aggression and suicide. Neuroscience. 2013;236: 160–85. Available: doi: 10.1016/j.neuroscience.2013.01.015

40. Perenyi A, Forlano R. Suicide in schizophrenia. Neuropsychopharmacol Hung. 2005;7: 107–17. Available: doi: 10.1586/ern.10.82

41. Bozorgmehr, A.1. Bozorgmehr A, Ghadirivasfi M, Tavakoli M, Rahmani H, Heydari F SAEI analysis of the genetic basis of suicidal behavior. PG 2018;28: 31–37. Ghadirivasfi M, Tavakoli M, Rahmani H, Heydari F, Shahsavand Ananloo E. Integrated analysis of the genetic basis of suicidal behavior. Psychiatr Genet. 2018;28: 31–37. Available: doi: 10.1097/YPG.0000000000000191.

42. Lee YJ, Kim S, Gwak AR, Kim SJ, Kang S-G, Na K-S, et al. Decreased regional gray matter volume in suicide attempters compared to suicide non-attempters with major depressive disorders. Compr Psychiatry. 2016;67: 59–65. Available: doi: 10.1016/j.comppsych.2016.02.013

43. Miller AB, McLaughlin KA, Busso DS, Brueck S, Peverill M, Sheridan MA. Neural correlates of emotion regulation and adolescent suicidal ideation. Biol Psychiatry Cogn Neurosci Neuroimaging. 2018;3: 125–132. Available: 10.1016/J.BPSC.2017.08.008

44. Lidow MS. Calcium signaling dysfunction in schizophrenia: a unifying approach. Brain Res Brain Res Rev. 2003;43: 70–84. Available: doi: 10.1016/S0165-0173(03)00203-0

45. Glausier JR, Lewis DA. Mapping pathologic circuitry in schizophrenia. Handbook of clinical neurology. 2018. pp. 389–417. doi:10.1016/B978-0-444-63639-3.00025-6

46. Broadbelt K, Jones L. Evidence of altered calmodulin immunoreactivity in areas 9 and 32 of schizophrenic prefrontal cortex. J Psychiatr Res. 2008;42: 612–621. doi:10.1016/j.jpsychires.2007.07.006

47. Saia-Cereda VM, Santana AG, Schmitt A, Falkai P, Martins-de-Souza D. The nuclear proteome of white and gray matter from schizophrenia postmortem brains. Mol Neuropsychiatry. 2017;3: 37–52. doi: 10.1159/000477299

48. Chin D, Means AR. Calmodulin: a prototypical calcium sensor. Trends Cell Biol. 2000;10: 322–8. Available: doi: 10.1016/s0962-8924(00)01800-6

49. Solà C, Barrón S, Tusell JM, Serratosa J. The Ca2+/calmodulin system in neuronal hyperexcitability. Int J Biochem Cell Biol. 2001;33: 439–55. Available: doi: 10.1016/s1357-2725(01)00030-9

50. Sharma RK, Parameswaran S. Calmodulin-binding proteins: A journey of 40 years. Cell Calcium. 2018;75: 89–100. Available: https://doi.org/10.1016/j.ceca.2018.09.002

51. Luo X-J, Li M, Huang L, Steinberg S, Mattheisen M, Liang G, et al. Convergent lines of evidence support CAMKK2 as a schizophrenia susceptibility gene. Mol Psychiatry. 2014;19: 774–83. Available: doi: 10.1038/mp.2013.103

52. Takemoto-Kimura S, Suzuki K, Horigane SI, Kamijo S, Inoue M, Sakamoto M, et al. Calmodulin kinases: essential regulators in health and disease. J Neurochem. 2017;141: 808–818. Available: doi: 10.1111/jnc.14020.

53. Means AR. Commentary: the year in basic science: calmodulin kinase cascades. Mol Endocrinol. 2008;22: 2759–2765. Available: doi: 10.1210/me.2008-0312

54. Fatemi SH, Folsom TD. The neurodevelopmental hypothesis of schizophrenia, revisited. Schizophr Bull. 2009;35: 528–548. Available: doi: 10.1093/schbul/sbn187

55. Zhang W, Wang D, Volk E, Bellen HJ, Hiesinger PR, Quiocho FA. V-ATPase V0 sector subunit A1 in neurons is a target of calmodulin. J Biol Chem. 2008;283: 294–300. Available: doi: 10.1074/jbc.M708058200

56. Schwaller B, Meyer M, Schiffmann S. “New” functions for “old” proteins: The role of the calcium-binding proteins calbindin D-28k, calretinin and parvalbumin, in cerebellar physiology. Studies with knockout mice. The Cerebellum. 2002;1: 241–258. Available: doi: 10.1080/147342202320883551

57. Schwaller B. The regulation of a cell’s Ca2+ signaling toolkit: The Ca2+ homeostasome. Springer, Dordrecht; 2012. pp. 1–25. doi:10.1007/978-94-007-2888-2_1

58. Permyakov EA, Uversky VN, Permyakov SE. Parvalbumin as a pleomorphic protein. Curr Protein Pept Sci. 2017;18: 780–794. Available: doi: 10.2174/1389203717666161213115746

59. Bastianelli E. Distribution of calcium-binding proteins in the cerebellum. The Cerebellum. 2003;2: 242–262. Available: doi: 10.1080/14734220310022289

60. Bullock WM, Cardon K, Bustillo J, Roberts RC, Perrone-Bizzozero NI. Altered expression of genes involved in GABAergic transmission and neuromodulation of granule cell activity in the cerebellum of schizophrenia patients. Am J Psychiatry. 2008;165: 1594–1603. Available: doi: 10.1176/appi.ajp.2008.07121845

61. Maloku E, Covelo IR, Hanbauer I, Guidotti A, Kadriu B, Hu Q, et al. Lower number of cerebellar Purkinje neurons in psychosis is associated with reduced reelin expression. Proc Natl Acad Sci. 2010;107: 4407–4411. Available: doi: 10.1073/pnas.0914483107

62. Brisch R, Bielau H, Saniotis A, Wolf R, Bogerts B, Krell D, et al. Calretinin and parvalbumin in schizophrenia and affective disorders: a mini-review, a perspective on the evolutionary role of calretinin in schizophrenia, and a preliminary post-mortem study of calretinin in the septal nuclei. Front Cell Neurosci. 2015;9: 393. Available: doi: 10.3389/fncel.2015.00393

63. Boley AM, Perez SM, Lodge DJ. A fundamental role for hippocampal parvalbumin in the dopamine hyperfunction associated with schizophrenia. Schizophr Res. 2014;157: 238–243. Available: doi: 10.1016/j.schres.2014.05.005

64. Abdul-Monim Z, Neill JC, Reynolds GP. Sub-chronic psychotomimetic phencyclidine induces deficits in reversal learning and alterations in parvalbumin-immunoreactive expression in the rat. J Psychopharmacol. 2007;21: 198–205. Available: doi: 10.1177/0269881107067097

65. Cunningham MO, Hunt J, Middleton S, LeBeau FEN, Gillies MJ, Gillies MG, et al. Region-specific reduction in entorhinal gamma oscillations and parvalbumin-immunoreactive neurons in animal models of psychiatric illness. J Neurosci. 2006;26: 2767–2776. Available: doi: 10.1523/JNEUROSCI.5054-05.2006

66. Piyabhan P, Tingpej P, Duansak N. Effect of pre- and post-treatment with Bacopa monnieri (Brahmi) on phencyclidine-induced disruptions in object recognition memory and cerebral calbindin, parvalbumin, and calretinin immunoreactivity in rats. Neuropsychiatr Dis Treat. 2019;15: 1103–1117. Available: doi: 10.2147/NDT.S193222

67. Servais L, Bearzatto B, Schwaller B, Dumont M, De Saedeleer C, Dan B, et al. Mono- and dual-frequency fast cerebellar oscillation in mice lacking parvalbumin and/or calbindin D-28k. Eur J Neurosci. 2005;22: 861–870. Available: doi: 10.1111/j.1460-9568.2005.04275.x

68. Farrecastany M, Schwaller B, Gregory P, Barski J, Mariethoz C, Eriksson J, et al. Differences in locomotor behavior revealed in mice deficient for the calcium-binding proteins parvalbumin, calbindin D-28k or both. Behav Brain Res. 2007;178: 250–261. Available: doi: 10.1016/j.bbr.2007.01.002

69. Wöhr M, Orduz D, Gregory P, Moreno H, Khan U, Vörckel KJ, et al. Lack of parvalbumin in mice leads to behavioral deficits relevant to all human autism core symptoms and related neural morphofunctional abnormalities. Transl Psychiatry. 2015;5: e525–e525. Available: doi: 10.1038/tp.2015.19.

70. Brown JA, Ramikie TS, Schmidt MJ, Báldi R, Garbett K, Everheart MG, et al. Inhibition of parvalbumin-expressing interneurons results in complex behavioral changes. Mol Psychiatry. 2015;20: 1499–1507. Available: doi: 10.1038/mp.2014.192.

71. Lungu O, Barakat M, Laventure S, Debas K, Proulx S, Luck D, et al. The incidence and nature of cerebellar findings in schizophrenia: A quantitative review of fMRI literature. Schizophr Bull. 2013;39: 797–806. Available: doi: 10.1093/schbul/sbr193.

72. Zhuo C, Wang C, Wang L, Guo X, Xu Q, Liu Y, et al. Altered resting-state functional connectivity of the cerebellum in schizophrenia. Brain Imaging Behav. 2018;12: 383–389. Available: doi: 10.1007/s11682-017-9704-0.

73. Behan ÁT, Byrne C, Dunn MJ, Cagney G, Cotter DR. Proteomic analysis of membrane microdomain-associated proteins in the dorsolateral prefrontal cortex in schizophrenia and bipolar disorder reveals alterations in LAMP, STXBP1 and BASP1 protein expression. Mol Psychiatry. 2009;14: 601–613. Available: doi: 10.1038/mp.2008.7

74. Martins-De-Souza D, Gattaz WF, Schmitt A, Rewerts C, Maccarrone G, Dias-Neto E, et al. Prefrontal cortex shotgun proteome analysis reveals altered calcium homeostasis and immune system imbalance in schizophrenia. Eur Arch Psychiatry Clin Neurosci. 2009;259: 151–163. Available: doi: 10.1007/s00406-008-0847-2

75. Martins-de-Souza D, Gattaz WF, Schmitt A, Maccarrone G, Hunyadi-Gulyás E, Eberlin MN, et al. Proteomic analysis of dorsolateral prefrontal cortex indicates the involvement of cytoskeleton, oligodendrocyte, energy metabolism and new potential markers in schizophrenia. J Psychiatr Res. 2009;43: 978–986. Available: doi: 10.1016/j.jpsychires.2008

76. Pinacho R, Villalmanzo N, Meana JJ, Ferrer I, Berengueras A, Haro JM, et al. Altered CSNK1E, FABP4 and NEFH protein levels in the dorsolateral prefrontal cortex in schizophrenia. Schizophr Res. 2016;177: 88–97. Available: doi: 10.1016/j.schres.2016.04.050

77. Velásquez E, Nogueira FCS, Velásquez I, Schmitt A, Falkai P, Domont GB, et al. Synaptosomal proteome of the orbitofrontal cortex from schizophrenia patients using quantitative label-free and iTRAQ-based shotgun proteomics. J Proteome Res. 2017;16: 4481–4494. Available: doi: 10.1021/acs.jproteome.7b00422

